# A visualizable hepatitis A virus and hepatitis C virus coinfection model *in vitro*: coexistence of two hepatic viruses under limited competition in viral RNA synthesis

**DOI:** 10.1101/709071

**Authors:** Wang Jiang, Pengjuan Ma, Gang Long

## Abstract

Hepatitis A virus (HAV) and hepatitis C virus (HCV) coinfection in patients usually leads to HCV suppression or clearance. Whether this suppression/clearance is caused by direct virus-virus interaction or indirect interactions is still unknown. Here, we present a robust and visualizable HAV/HCV coinfection model for investigating their interactions *in vitro*. We find that HAV super infects HCV-persistently-infected Huh-7.5.1 cells without obvious virus-virus exclusion and vice versa, while there is a mild reciprocal viral interference in coinfection. Through single-cell scale confocal microscopy analysis and treating co-infected cells with rNTPs or antivirals e.g. Sofosbuvir, Simeprevir, and Ledipasvir, we find that HAV and HCV exploit cellular machinery in a compatible manner, but with reciprocal competition for rNTPs in their RNA synthesis. In conclusion, our findings reveal the absence of direct HAV-HCV interaction but the presence of indirect interaction, which may be due to limited competition in viral RNA synthesis. Therefore, we propose that suppression/clearance of HCV in HAV/HCV coinfected patients is probably due to indirect interactions e.g. viral competition or immunological interactions.

**Author summary:** Virus-virus interactions could usually be categorized in three aspects: (1) direct interaction induced by interaction of viral gene or gene products; (2) indirect interaction mediated by immune response or (3) host environmental alteration. Hepatitis A virus (HAV) and hepatitis C virus (HCV) are two important causatives of human hepatitis. In patients, their coinfection often induces HCV suppression/clearance, and sometimes may lead to fulminant hepatitis. In this study, we find that a limited reciprocal viral interference occurs in HAV/HCV coinfection *in vitro*, which results from a viral competition of rNTPs in viral RNA synthesis. Our study provides a new paradigm of studying virus-virus interactions in single-cell scale. To our knowledge, we identify the rNTPs competition in virus coinfection for the first time by experiment, though several mathematical modelings have predicted the resource competition in virus coinfection.

## Introduction

Hepatitis A virus (HAV) and hepatitis C virus (HCV) are both hepatotropic viruses, with HAV usually causing acute infection and HCV often resulting in chronic infection [1]. HAV is a food-borne virus, which is epidemic in hygiene-poor regions and often causes a sporadic outbreak in developed countries [2]. Acute HAV infection often induces nausea, vomiting, diarrhea, jaundice or even liver failure [2]. Although HAV vaccine was accessible two decades ago [3], potent antivirals against HAV is still in demand [4]. In contrast, HCV is a blood-borne virus, chronically infecting approximately 80 million people worldwide and often leading to liver cirrhosis and even hepatocellular carcinoma [5]. Although efficient direct-acting antivirals (DAAs) against HCV dramatically improved the therapy of chronic hepatitis C, the HCV vaccine has not yet been available [5].

HAV and HCV are both single-stranded positive-sense RNA viruses, belonging to the Picornaviridea and Flaviviridae respectively [6]. They exhibit many similar biological features. For example, (1) their robust replication *in vitro* are induced by the cell-culture adapted mutations under experimental selection [7, 8]; (2) They have a comparable genome structure, which encodes a single polyprotein that is processed by their viral or cellular proteases into separate viral proteins [9, 10]; (3) Their infections in cells alter the morphology of endoplasmic reticulum (ER). e.g. Tubular-vesicular network by HAV [11] versus membraneous web by HCV [12]; (4) They evade the innate immunity via the same strategy: cleaving IPS-1 to inactivate RIG-I antiviral pathway [13, 14]. On the other hand, differences between HAV and HCV are obvious. For example, (1) HCV has its own viral membrane proteins E1 and E2 [9], while HAV is naked or quasi-enveloped through hijacking cellular membranes [15, 16]; (2) It is well-known that HCV utilizes many cellular receptors for its entry [9], whereas HAV entry step is still enigmatic as the only known receptor—TIM1 seems not essential for HAV infection [17].

HAV infection sometimes leads to severe or even fulminant hepatitis in the patient with chronic hepatitis C [18], which is disputable in different retrospective studies [19]. In contrast, it is a common phenomenon that HAV infection leads to HCV suppression or clearance in HAV/HCV coinfected patients [20, 21]. Indirect interaction mediated by immune response might be one reason for HCV suppression/clearance. e.g. Cacopardo et al. demonstrated that HAV superinfection-induced IFN-γ production was associated with HCV clearance [22]. Nevertheless, whether the direct virus-virus interaction or some other indirect interactions could contribute to HCV suppression/clearance is still unknown.

In this study, we established a robust and visualizable HAV/HCV coinfection model in Huh-7.5.1 cell to investigate HAV-HCV interactions. Our coinfection model is suitable for studying virus-virus interactions without considering immunological interaction, as Huh-7.5.1 cell is immunocompromised in response to virus infection [23, 24]. Our data revealed the absence of direct HAV-HCV interaction but the presence of reciprocal competition in viral RNA synthesis.

## Results

### The absence of HAV-HCV exclusion in HAV superinfection revealed by a dual fluorescence reporter system

For visualizing real-time HAV/HCV coinfection with convenience, we combined two real-time detecting systems of HAV and HCV infection in Huh-7.5.1 [25, 26]. In brief, for HAV detection, C508 at IPS-1 was mutated to R508R for excluding the HCV NS3/3A cleavage and GFP (green fluorescent protein) was used (Fig 1A); For HCV detection, RFP (red fluorescent protein) was used (Fig 1A). Because HAV 3ABC and HCV NS3/4A cleave IPS-1 at Q427 and C508 (Fig 1A), relocalization of GFP or RFP from cytosol to nucleus could report HAV or HCV infection in the same cell without disturbance (Fig1A, S1 FigC). We designated this dual fluorescence reporter system as Huh-7.5.1-GA/RC, which supported robust infection of both HAV and HCV (S1 Fig A-B). HAV superinfection in the patient with chronic hepatitis C is a common phenomenon, we first investigated the HAV superinfection in HCV persistently infected Huh-7.5.1-GA/RC cells (Fig 1B). To note, we applied a high MOI of HAV and HCV infection for ensuring that 90% cells were HAV/HCV co-infected. Seven days post HCV infection, HCV succeeded to infect almost 90% cells (Fig 1C) and HCV infection did not disturb TIM-1 expression (S1 Fig D), after which we initiated HAV superinfection. We observed that at least 90% cells were HAV/HCV coinfected from the time of 48 hours post HAV infection to the end of our experiment (Fig 1D). These data demonstrated that HAV superinfection did not induce obvious exclusion of HCV replication in the immunocompromised model and there was no obvious superinfection exclusion.

**Fig 1.**
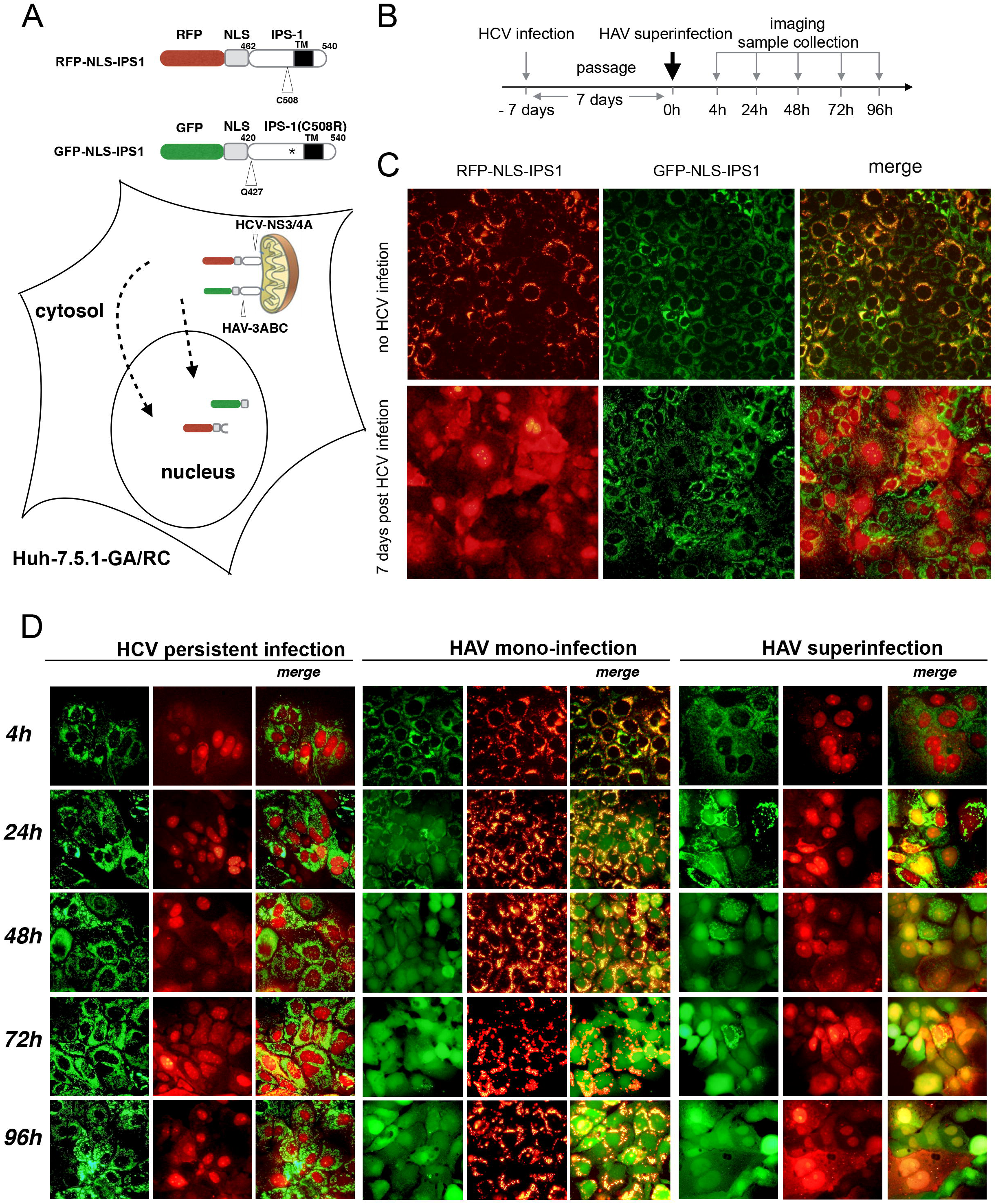
A visualizable HAV/HCV coinfection model revealed the absence of virus-virus exclusion in HAV superinfection. (A) An IPS-1-based reporter system for dual detection of HAV and HCV. C-terminal transmembrane domain(TM) of IPS-1 directs IPS-1 to the outer membrane of mitochondria. Nuclear localization signal(NLS) directs the protein to the nucleus. The HCV NS3/4A protease cleaves IPS-1 at C508 (arrow indicated) and HAV 3ABC protease at E427 (arrow indicated). asterisk indicates the mutation site C508R. (B) Timeline of HAV superinfection assay. 7 days before HAV superinfection, Huh-7.5.1-GA/RC cells were seeded on 6-well plate and inoculated with HCV virus stock (MOI≈2.5) the next day. After 5 days, the HCV-infected Huh-7.5.1-GA/RC cells were seeded on 24-well plates for the next day’s HAV superinfection. At the indicated time point, images were captured and cells were harvested. (C) More than 90% of cells were HCV-infected 7 days post HCV infection. (D) Real-time visualization of HAV superinfection. Total magnification of photographs is 200X power. Data represent one of three independent assays.

### Reciprocal viral interference between HAV and HCV in HAV superinfection

In order to confirm the above finding revealed by the reporter system, we conducted a comprehensive analysis of HAV superinfection via RT-qPCR, western blot, and infectivity titration. HAV cytopathogenic and non-cytopathogenic strain HM175/18f and HM175/p16 were used, with the former replicating more robustly than the latter (Fig 1A-C). Compared to their mono-infection, viral proteins expression of HAV and HCV reduced slightly in superinfected cells (Fig 2A, Fig 2D). Consistent with this, by RT-qPCR, we observed a mild decrease (5-fold ∼ 10-fold reduction) of both HAV and HCV intracellular RNA copies in HAV superinfection (Fig 2B, Fig 2E). In addition, we found that extracellular HAV infectivity and VP2 decreased approximately 10-fold in HAV superinfection compared to its mono-infection, whereas extracellular HCV infectivity and core were not disturbed (Fig 2C, Fig 2F, S2 Fig A-B).

**Fig 2.**
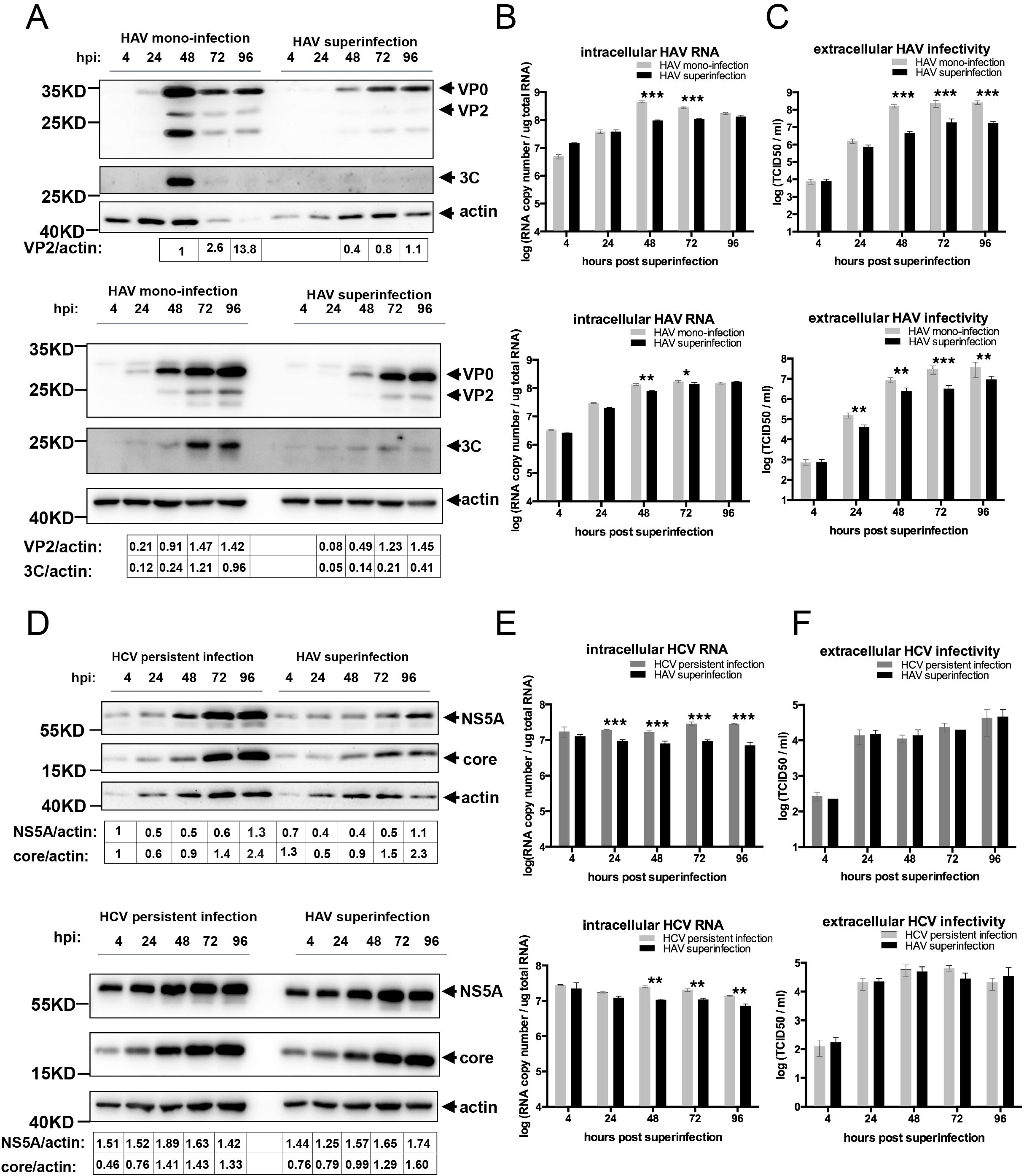
A reciprocal mild interference between HAV and HCV in HAV superinfection. Cells harvested in Fig 1 were further analyzed by Western blot, RT-qPCR and infectivity titration. (A)(D) Western blot analysis of viral protein expression. Band intensity was estimated by Fiji and normalized for actin. (B)(E) Quantification of intracellular viral RNA by RT-qPCR. The results were normalized by quantification of cellular total RNA. (C)(F) Infectivity analysis of extracellular viruses by TCID50. hpi represents ‘hours post infection’. Upper panel represents the result of HM175/18f superinfection. Down panel represents the result of HM175/p16 superinfection. Data represent the mean of three independent assays.

It is well-known that HAV/HCV infection could lead to cell apoptosis *in vitro*, which was confirmed by Annexin-V / PI staining in our study (S3 Fig B). In addition, we found that the percentage of HAV-induced cell loss and apoptosis was less in HAV superinfection than in HAV mono-infection (S3 Fig A-B), which suggested that the attenuation of HAV replication by HCV may lessen the cell death. We next determined whether the attenuation of HCV replication result from HAV-induced cell death. However, low MOI HM175/18f superinfection and long-term HM175/p16 superinfection still induced a reciprocal interference of their RNA synthesis (S4 Fig A-D). These data suggested that the attenuation of HAV/HCV replication in their coinfection should not result from cell death.

Direct virus-virus interaction usually exhibits dramatic inhibition of another virus during coinfection. e.g. over-expression of Borna disease virus (BDV) nucleocapsid components prevented a subsequent infection of a different BDV strain [27]. In this HAV/HCV coinfection model, HAV and HCV replications in Huh-7.5.1 cell were robust, which would provide enough amount of viral proteins for direct interactions. Besides, over-expression of HCV viral structural protein E1, E2, and core in Huh-7.5.1 cells did not affect HAV infection (S5 Fig A-C). However, we just observed a limited reciprocal viral interference between HAV and HCV (Fig 2), which suggested that direct interactions between HAV and HCV were absent in HAV/HCV coinfection.

### Neighbored HAV/HCV replication complexes and competition in their RNA synthesis

Next, we dedicated to investigating the mechanism of their reciprocal limited viral interference by confocal microscopy. The localization of their replication complexes (RCs) was observed by labeling their negative-sense RNA by FISH (fluorescent in-situ hybridization). In HAV/HCV coinfected single-cell, their RCs did not co-localize in the cytosol (Fig 3A) as the Pearson’s correlation coefficient of HAV/HCV negative RNA is close to 0 (Fig 3C). Moreover, by 3D image reconstruction, we found that their RCs were neighbored to each other (Fig 3B). Besides, we simulated viral RCs via RNA spots detected by FISH, which were represented as balls (Fig 3D). To note, balls were rendered by the same rule that minimum radius of one ball was determined automatically by iMaris in order to cover one separate RNA spot signal. We found that co-infected cell and mono-infected cell owned equivalent numbers of HAV/HCV RNA spots (Fig 3E), whereas the signal intensity of RNA spots slightly reduced in the coinfected cell (Fig 3F). Furthermore, strand-specific RT-qPCR confirmed both decrease of their negative- and positive-sense RNAs in HAV/HCV coinfection (S6 Fig). These data suggested that HAV/HCV coinfection did not alter space occupation of individual viral RCs, but may lead to attenuation of their RNA synthesis.

**Fig 3.**
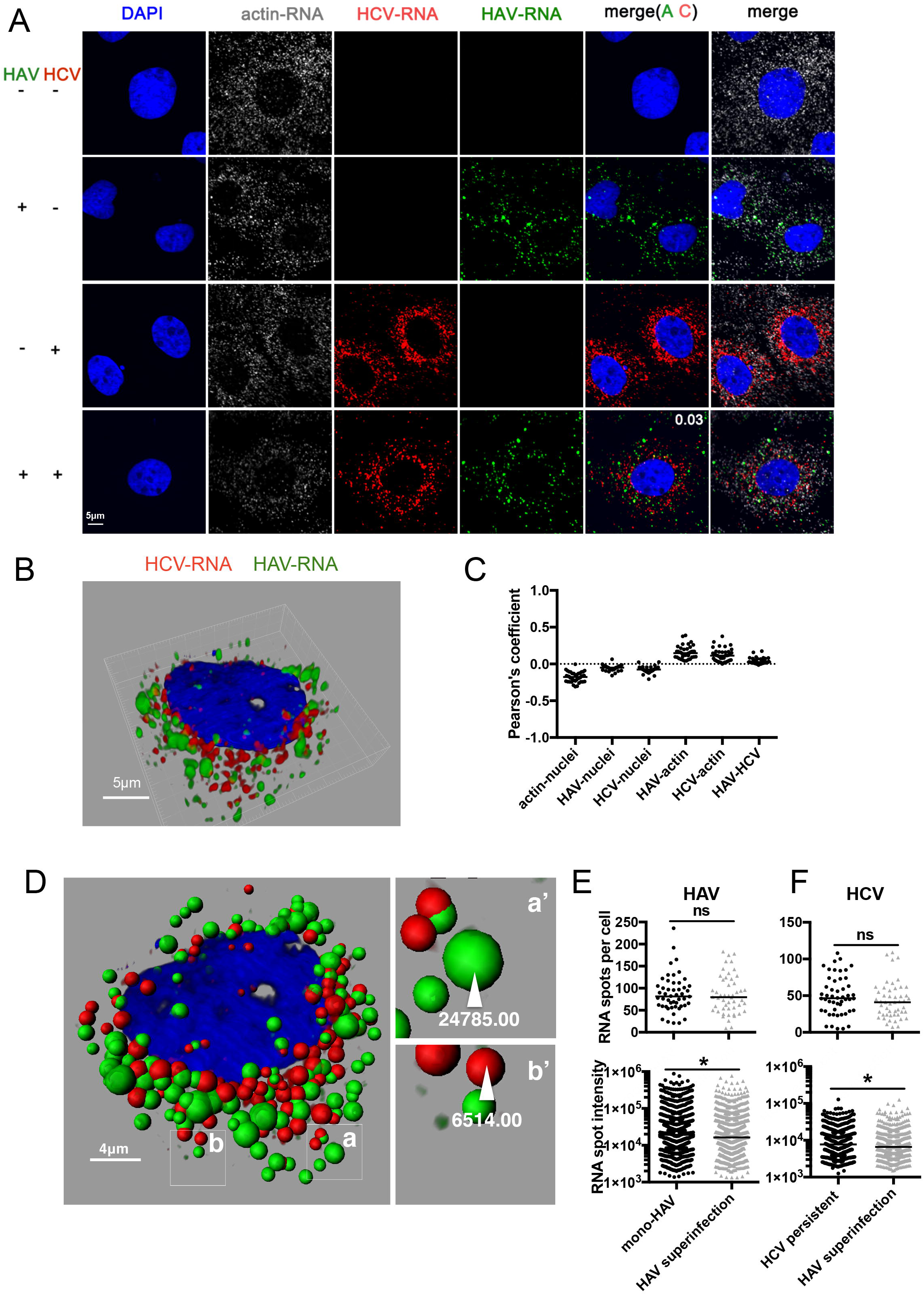
HAV/HCV RCs in the co-infected cell were neighbored and competition in their RNA synthesis. Forty-eight hours post HAV (HM175/18f) superinfection, cells were fixed by 4% PFA and analyzed by FISH as described in methods. Pseudo-colors were applied using FV10-ASW. (A) Virus negative-sense RNA ((-)RNA) observed by FISH. A white number in pictures represents the Pearson’s correlation coefficient between HAV (-) RNA and HCV (-)RNA, which was calculated by FV10-ASW. (B) 3D-representation of HAV and HCV RCs in the HAV-superinfected cell. Image stacks were first deconvolved using AutoQuant X3 and then rendered into a 3D image using Imaris as described in methods. (C) Pearson’s correlation coefficient analysis of two analyzed elements in co-infected cells as shown in the figure. At least 40 HAV-superinfected-cells were analyzed. The correlation between virus (-)RNA and actin mRNA was used as a negative control. (D) An example of RNA spot simulation into the ball in HAV and HCV co-infected Huh-7.5.1 cell. The number of RNA balls was calculated in at least 30 cells. Total intensity in one spot was summed to a value by iMaris as shown in (a’) and (b’). (E) Upper: Numbers of HAV RNA spots in single-cell. Down: Intensity in one HAV RNA spot. (F) Upper: Numbers of HCV RNA spots in single-cell. Down: Intensity in one HCV RNA spot.

### rNTPs treatment to HAV/HCV coinfection alleviated viral competition in viral RNA synthesis

Both HAV and HCV synthesize their RNA template using rNTPs via their viral RNA-dependent RNA polymerases (Rdrp). Considering the above imaging data, we hypothesized that rNTPs competition in RNA synthesis may result in the reciprocal limited viral interference in HAV/HCV coinfection. Then, we inoculated cells with rNTP four hours post HAV superinfection to see whether extra-added rNTP could rescue the attenuation of their replication in HAV/HCV coinfection. As expected, rNTP increased both HAV and HCV replication significantly in coinfection (Fig 4A, Fig 4B). In addition, rNTP increased HAV replication in HAV mono-infection, while it has no effect on HCV replication in HCV persistent infection (Fig 4A, Fig 4B). Previously, we found that Sofosbuvir—an analog of rNTPs, inhibited HAV replication with IC50 of 6.8μM, while it inhibited HCV replication more efficiently with IC50 of 0.05μM [26]. Here, in HAV/HCV coinfection, antiviral activity of sofosbuvir against HAV and HCV became worse, compared to that in their mono-infections (Fig 4C, Fig 4D). Especially when the Sofosbuvir concentration was at 2000nM, inhibitory effect on HAV was disappeared, though at the same concentration in HAV mono infection Sofosbuvir inhibited approximately 50% HAV replication (Fig 4C). The antiviral activity of Sofosbuvir against RNA virus is to inhibit viral Rdrps via binding to it, which suggested that HAV and HCV Rdrps even compete for Sofosbuvir. From this result, we demonstrated that rNTPs should be a crucial factor in HAV/HCV coinfection, for which HAV and HCV Rdrps strive against each other for their RNA synthesis.

**Fig 4.**
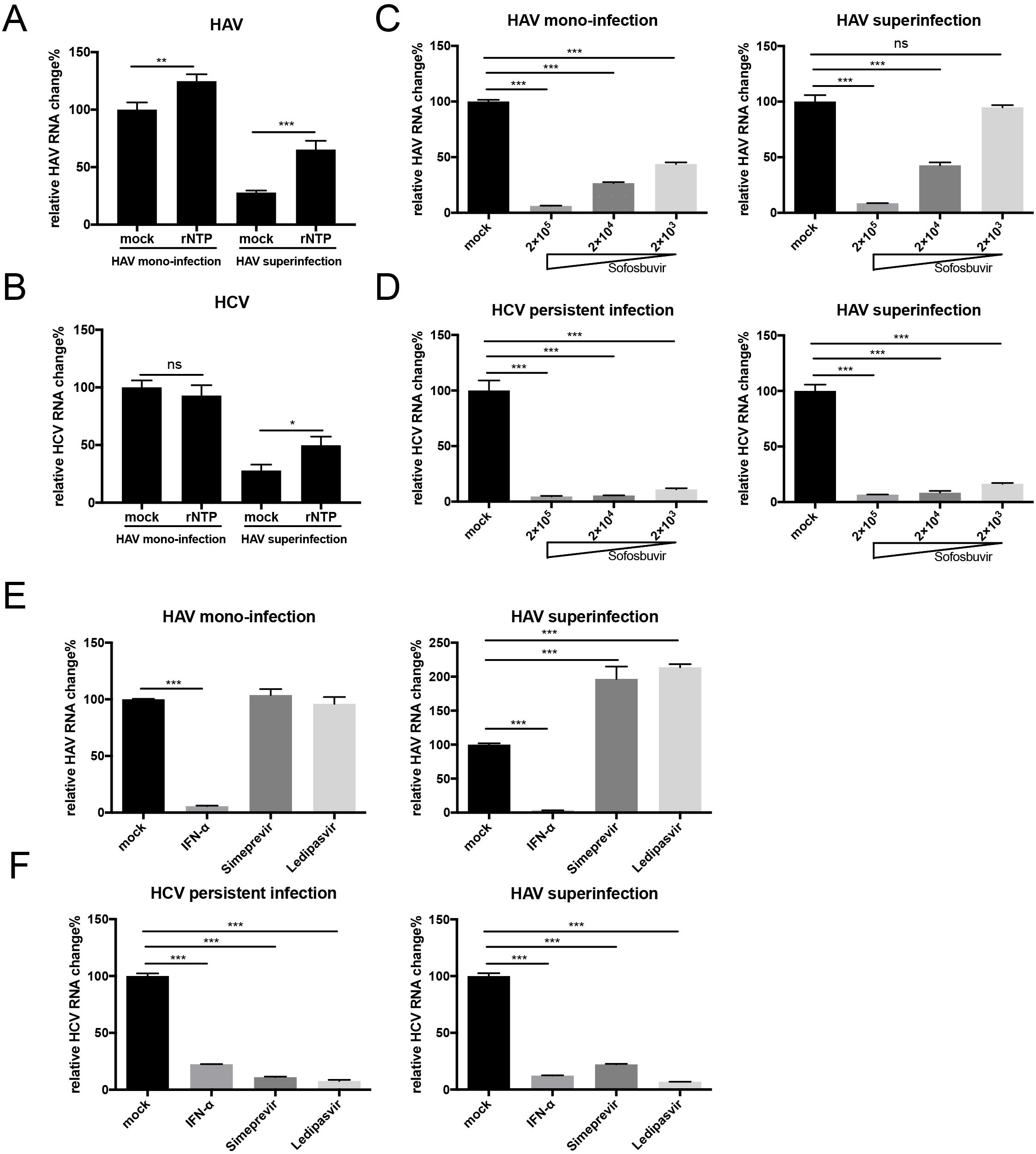
Extra-added rNTPs increases both HAV and HCV replication in coinfection. rNTPs and other drugs were added at the time of four hours post HAV(HM175/18f) infection. The working concentrations of rNTP, IFN-α, Simeprevir, and Ledipasvir were 0.1mM, 1000IU/ml, 200nM and 200nM respectively; Sofosbuvir concentration (nM) was used as indicated. Mock represents 0.2% DMSO. 48 hours after HAV infection, cells were washed once with 1×PBS and harvested. RT-qPCR was used to quantify the intracellular viral RNA amounts and relative change was created by dividing the mock. (A) Effect of rNTPs on HAV replication and (B) Effect of rNTPs on HCV replication. (C) Effect of sofosbuvir on HAV replication and (D) Effect of sofosbuvir on HAV replication. (E) Effect of indicated drugs on HAV replication. (F) Effect of indicated drugs on HCV replication. Data represent the mean of three independent assays.

### HCV DAAs treatment to HAV/HCV coinfection lessened the attenuation of HAV replication

On the other hand, Simeprevir and Ledipasvir, which specifically inhibited HCV replication, increased HAV replication in HAV/HCV coinfection (Fig 4E, Fig 4F). Finally, we found that IFN-α inhibited both viruses replication (Fig 4E, Fig 4F). These results indicated that the attenuation of HAV replication is due to the competition from HCV replication.

### HAV/HCV assembly and releasing were not overlapped

By immunofluorescence analysis (IF) of their structural protein, we found that HCV core and HAV VP3 were neighbored but did not show any significant overlapping (the median of their Pearson’s correlation coefficient is 0.144) (Fig 5A, Fig 5C). In addition, we found the colocalization of HCV core and lipid droplets (LDs), which is consistent with that HCV assembly depends on the lipid pathway [28]. However, there was no colocalization between HAV VP3 and LDs (Fig 5B, Fig 5C). Considering eHAV (enveloped HAV) was shown to acquire the exosome-associated membrane [15, 16] and HCV are reported to be tightly associated with exosome pathway [29], we asked whether HAV could hijack HCV membrane or not (Fig 5D). Then, we did a pull-down analysis of HCV virion by immunoprecipitating E2 in cell culture, to check if HAV can be co-precipitated or not. Approximately 50% HCV infectivity or genomic RNA was pull downed in the eluate from both HAV superinfection and HCV persistent infection. However, we can not observe detectable HAV infectivity or its genomic RNA co-precipitated from HCV virions (Fig 5E, Fig 5F). These results indicated that the HAV/HCV assembly and release should not be overlapped.

**Fig 5.**
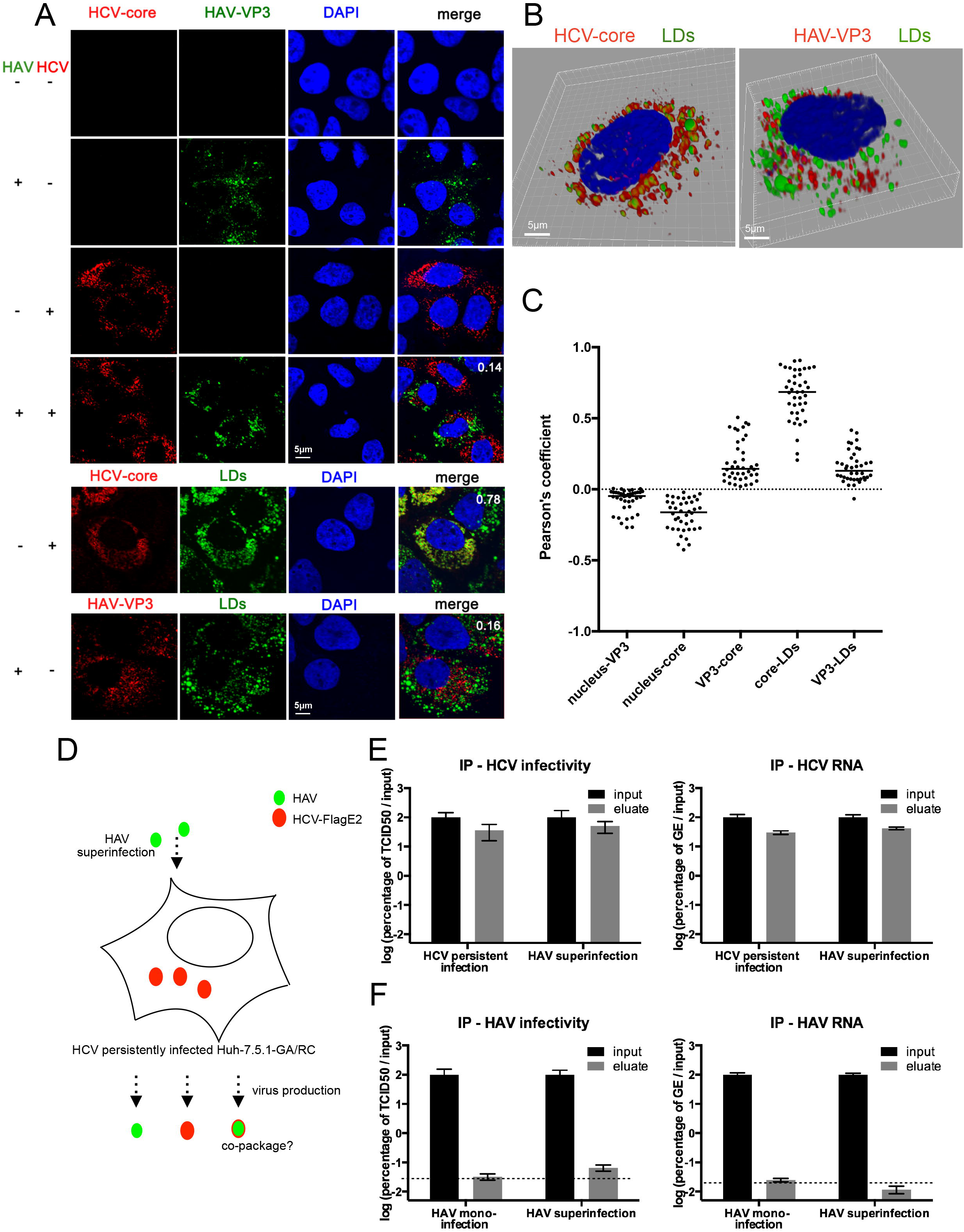
HCV and HAV assembly and releasing was not overlapping. (A) Immunofluorescence (IF) analysis of viral structural proteins and the lipid droplets (LDs) by confocal microscopy. The white number represents the Pearson’s correlation coefficient. (B) 3D image of viral protein and lipid droplets in HAV mono-infected (left) and HCV mono-infected (right) Huh-7.5.1 cell. (C) Pearson’s correlation coefficient between the two analyzed elements. The correlation of viral structural protein and nucleus were used as a negative control. At least 40 HAV-superinfected-cells were analyzed. (D) Schematic diagram of the speculated co-packaging in superinfection setup. Immunoprecipitation (IP) assay was applied to pull-down JC1E2flag particles with anti-flag beads. Viral genomes and infectivity were quantified by RT-qPCR and TICD50 assay. Cell cultures from mono-HAV infected cells and HCV-persistently-infected cells were used as a negative and positive control. (E) HCV infectivity and genome (GE) RNA were shown as the percentage of the input. (F) HAV infectivity and genome (GE) RNA were shown as the percentage of the input. The dashed line indicates the detection limit. Data represent one of three independent assays.

### Reciprocal viral interference between HAV and HCV in HCV superinfection

As limited competition was revealed in the HAV superinfection model without direct viral interaction, we hypothesized that HAV and HCV should co-exist with limited interference whatever the order of superinfection. So we conducted HCV superinfection in HAV infected cells and ensured 90% cells were coinfected (data not shown). As expected, their RNA synthesis, protein expression, and virus production showed reciprocal limited interference (Fig 6).

**Fig 6.**
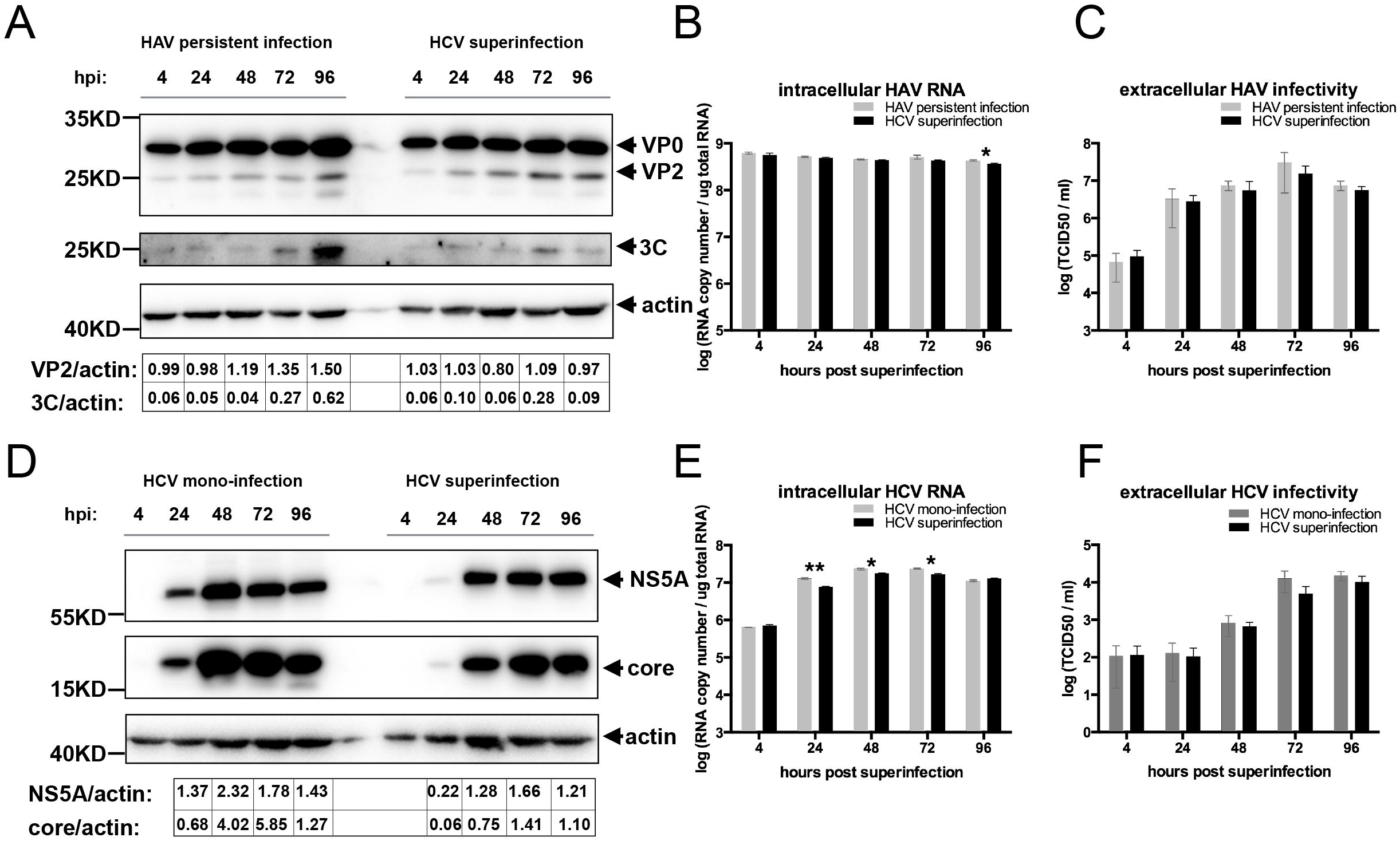
A reciprocal mild interference between HAV (HM175/p16) and HCV in HCV superinfection. HCV superinfection was initiated after persistent HAV infection establishes in Huh-7.5.1-GA/RC cell. (A)(D) Western blot analysis of viral protein expression. Band intensity was estimated by Fiji and normalized for actin. (A) The HAV non-structural protein (3C) and structural protein (VP2) were detected. (D) The HCV non-structural protein (NS5A) and structural protein (core) were detected. (B)(E) Quantification of intracellular viral RNA by RT-qPCR. The results were normalized by quantification of cellular total RNA. (C)(F) Infectivity analysis of extracellular viruses by TCID50. Data represent one of three independent assays. hpi represents ‘hours post infection’.

## Discussion

In this study, we established a visualizable *in-vitro* HAV/HCV co-infection model and made a comprehensive analysis of HAV-HCV interaction in Huh-7.5.1 cell. We monitored the real-time process of HAV/HCV coinfection (including HAV and HCV superinfection) and found that there was no obvious virus-virus exclusion when more than 90% cells were HAV/HCV coinfected. However, a reciprocal limited viral interference was observed, which was consistent with a previous study that HAV and HCV replicases have limited competition in a sub-genomic replicon system [30]. In addition, we identified the mechanism of limited interference.

Virus-virus interactions are common phenomena in natural hosts and can be categorized into three aspects: (1) direct interactions of viral genes or gene product, (2) indirect interactions that result from alterations in the host environment, and (3) immunological effect [31]. First, we exclude the possibility of direct HAV-HCV interactions resulting in limited viral interference. Second, we found that viral competition of rNTP in their RNA synthesis resulted in the reciprocal interference in HAV/HCV coinfection. It is possible that the attenuation of RNA synthesis would affect the downstream of the viral life cycle such as translation and assembly. However, we can not exclude that the limited viral competition occurred throughout their virus life cycles in coinfection. For example, 1) mild reciprocal interference of HAV and HCV viral protein expression indicated that their RNA template might compete to recruit ribosomes for translation; 2) interference on HAV extracellular infectivity might be due to the competition of some host factors such as Rab27a [32], because HCV and HAV releasing are tightly related to the exosome pathways [16, 29, 33]. In our model, HAV replication capacity and secretion efficiency was higher than that of HCV, which suggested that HAV need more resource than HCV. However, HAV and HCV can not directly disturb each other i.e. HAV suffers more pressure from the competition than HCV. This may explain why the attenuation of HAV life cycle (especially HM175/18f) was more obvious in coinfection.

Resource competition in virus coinfection has been predicted in several mathematical modelings. e.g. Lubna Pinky et al. predicted in their study that rhinovirus, the fastest-growing virus, reduces replication of the remaining viruses during a coinfection, while parainfluenza virus, the slowest-growing virus is suppressed in the presence of other viruses [34]. To our knowledge, our study is the first experimental one to describe resource that the two virus compete for rNTPs in viral RNA synthesis.

In patients, HAV/HCV coinfection would lead to HCV suppression/clearance. In this *in-vitro* coinfection model, we did not detect any dramatic inhibition of HCV or direct HAV-HCV interactions, which indicated the absence of direct virus interaction in HAV/HCV coinfected patients. Both HAV and HCV are cell-culture adapted strains, which replicate robustly in Huh-7.5.1 cells. *In vivo*, the replication capacity of wild-type strains in infected patients should be an important factor that determines the occurrence of competition between HAV and HCV *in-vivo*. For example, HM175/p16 induces weaker viral interference than HM175/18f in HAV/HCV coinfection (with the former replicating less robustly than the latter). In addition, the competition also depends on the proportion of HAV-HCV coinfected hepatocytes in the patient’s liver, which is lack in clinical data. In coinfected patients, an indirect interaction mediated by immunological interaction may be another reason. Cacopardo et al. found that increased IFN-γ production, stimulated by HAV superinfection, is associated with the undetectable HCV RNA in patients [22]. Besides, HAV and HCV were reported to have an opposite effect on T-reg activation [35, 36]. Future study is needed to investigate whether HAV-induced T-reg inactivation or other immunological interaction could result in HCV suppression/clearance.

In conclusion, we report a visualizable cell culture model system allowing the study of HAV–HCV coinfection *in vitro*. Our data demonstrate the absence of direct interaction between these two viruses and indicate that indirect interaction such as viral competition may explain the HCV suppression/clearance observed in patients. Our findings provide deeper insights into the pathogenesis of HAV–HCV coinfection and open a new vision of studying two viruses interaction in single-cell level.

## Materials and methods

### Plasmids and antibodies

Plasmids pFK-JC1E2^flag^ have been previously described [37]. Plasmids pGEM3-HM175/18f was a kind gift from Stanley M. Lemon [15]. Plasmids pWPI-blr-GFP-NLS-IPS1, pWPI-puro-RFP-NLS-IPS1 for the fluorescent reporter system were constructed as described previously [26]. Briefly, fragments of GFP-NLS, RFP-NLS, NLS-IPS1(420/540), NLS-IPS1(462/540) were amplified by PCR. Then GFP-NLS-IPS1(420/540), RFP-NLS-IPS1(462/540) were amplified by overlapping PCR, and inserted into lentiviral plasmid pWPI. Primers are listed in S1 table. Mouse anti-NS5A, rabbit anti-NS5A, rabbit anti-core have been described previously [38]. Rabbit anti-3C, rabbit anti-VP2 were made in home. Other antibodies were purchased as follows: Mouse anti-VP3 (Lifespan), goat anti-actin (Santa-cruz), goat anti-rabbit-HRP (ZSGB-Bio), goat anti-mouse-HRP (ZSGB-Bio), goat anti-mouse/rabbit labeling with Alexa-fluor-488/555® (lifetechnologies). Sofosbuvir, Simeprevir, and Ledipasvir were purchased from MedChem Express. IFN-α was a gift from X.Liu. rNTPs were purchased from Promega.

### Cells and viruses

HEK293T, Huh-7.5.1, and its derivative Huh-7.5.1-GA/RC were cultured in DMEM medium supplemented with 2mM L-glutamine, nonessential amino acids, 100U penicillin per ml, 100μg streptomycin per ml and 10% FBS (complete DMEM). Production of JC1E2^flag^ and HM175/18f have been described previously [15, 38]. Virus stock was stored at −80□ after filtration with 0.45μm filters (Millipore). HM175/p16 virus was a kind gift from Stanley Lemon and Zongdi Feng. Infectivity of HAV and HCV were titrated by TCID50 as previously described [38]. Briefly, Huh-7.5.1 cells were seeded in 96-well plate, and one day later, the virus was added to the plate in serial dilution. 3 day later, cells were fixed using cold ethanol and hybridized with mouse anti-NS5A for HCV, while titration of HAV was determined by the HAV fluorescence reporter system—Huh-7.5.1-GA as described previously [26]. For generation of Huh-7.5.1-GA/RC reporter cells, lentiviruses carrying GFP-NLS-IPS1(420/540) and RFP-NLS-IPS1(462/540) were prepared as previously described [38]. Then Huh-7.5.1 was co-transfected by the lentiviruses and screened under blasticidin (4μg/ml) and puromycin (1μg/ml) to generate Huh-7.5.1-GA/RC cell line.

### Western blot analysis

Western blot was applied as previously described [38]. Briefly, Samples were collected in Laemmli buffer and separated by SDS polyacrylamide gel electrophoresis. Proteins were transferred onto PVDF membrane, which was blocked using 5% dried milk and incubated with indicated primary antibody and secondary antibody. Bands were visualized by ECL reagent (Bio-Rad) under Tanon 4200. Software Fiji was applied to estimate the band intensity.

### RT-qPCR assay for quantification of viral RNA

Cells were washed with 1×PBS once and the supernatant was filtered by 0.45μm filters prior to adding the TRNzol Universal Reagent (Tiangen). Total cellular RNA was extracted according to the manufacturer’s protocol (Tiangen) and the concentration of RNA was determined by NanoDrop 2000. Extracted RNA was reverse-transcribed into cDNA with FastQuant RT kit (Tiangen) and the cDNA were subjected to ABI 7900HT or ABI Q6 with SYBR Green (Tiangen) for qPCR. Serial dilution of the equivalent volume of the in-vitro transcribed viral genomic RNA was used to create a standard curve for converting Ct values into absolute gnomonic numbers. For strand-specific RT-qPCR, RT-primers adapted with a specific tag sequence, complementary to the region in 5’-UTR of HAV and HCV genome, were used to reverse transcription. Then tag-primer coupled with forward or reverse qPCR primer were used for qPCR. For acquiring minus-strand RNA template, cDNA of the negative strands of HAV/HCV 3’UTRs were cloned together in pCDNA3.1 (pCDNA3.1-AC minus) for in-vitro transcription. Primers for RT-qPCR and cloning pCDNA3.1-AC minus were listed in S2 table.

### Fluorescence imaging

For visualizing the real-time coinfection, Huh-7.5.1-GA/RC cells were observed and taken photos under Olympus IX53 fluorescent microscopy. For immunofluorescence (IF) of viral proteins, Huh-7.5.1 cells grown on glass coverslips in the 24-well plate were fixed with 4% paraformaldehyde for 15 minutes, permeabilized with 0.5% Triton-100, and blocked with 5% calf serum. Cell was then incubated with the primary antibodies for 1h at room temperature, washed, and incubated with Alexa fluor555® goat anti-rabbit or Alexa fluor 488® anti-mouse for 1h at room temperature. After washing, the cells were incubated with DAPI (beyotime) or Bidopy495/503 (lifetechnologies) for 5 minutes. Finally, glass coverslips were mounted on glass slides with Fluoromount Aqueous mounting medium (sigma). For fluorescent in situ hybridization (FISH) of viral (-)RNA, the QuantiGen® ViewRNA ISH cell assay kit (Affymetrix) was used according to the manufacturer’s protocol, except not adding protease. Briefly, cells were fixed with 4% paraformaldehyde for 30 minutes at room temperature, incubated with detergent solution for 5 minutes. Probe sets specific for viral (-) RNA were hybridized to the cells at 40□ for 3 hours, pre-amplifier and amplifier DNA probes were sequentially inoculated with cells at 40□, each for 30 minutes. Cells were then incubated with fluorescently labeled probes, specific to the amplifier DNA probes at 40□ for 30 minutes. DAPI dying was conducted at last and coverslips were mounted onto a glass slide with Fluoromount Aqueous mounting medium (Sigma). Images were captured under the Olympus FV-1200 laser-scanning confocal microscopy and filter sets are as follows: DAPI—408nm, HAV RNA—550nm, HCV RNA —650nm, actin RNA—488nm. Pearson’s correlation coefficient was calculated with the software FV10-ASM. For 3D reconstruction, one cell was scanned with approximately 8 optical slices along the Z-axis (0.63μm/slice) using confocal microscope FV-1200, then stacks of slices were processed by 3D blind deconvolution using AutoQuant X3. Analysis and projection of 3D images were used by Imaris. All images were captured and processed using the same parameter in order to compare RNA spot numbers and intensity in different cells.

### Immunoprecipitation for pulling down HCV particles

The cell culture medium was first condensed to 10% primitive volume by Amicon®100K Centrifugal Filter (Millipore),then were incubated with Flag-affinity gel (Sigma) at 4□ overnight.

After washing the gel with 1×PBS 10 times, Flag-peptide(Sigma) equal to 5 × bed volumes of Flag-affinity gel was used for eluting HCV particles. TCID50 and RT-qPCR were applied to measure viral infectivity and RNA copy number in input and eluate.

### Cell viability and apoptosis detection assay

Cells were seeded in 96-well plates at a density of 1.0 × 10□ cells/well and cultured overnight. At different time points post-infection, cell viability/proliferation was determined by the MTT assay [39]. For detection of apoptosis/necrosis, Annexin V-EGFP Apoptosis Detection Kit (BioVision) was used according to the manufacturer’s protocol. In brief, the culture supernatant was removed softly to avoid the loss of the apoptotic cell, then cells were incubated with 1×binding buffer dissolving Annexin V-EGFP and PI for 5 minutes. Images were captured immediately under Olympus IX53 fluorescent microscopy.

### Statistical analysis

For the time course assay, significance value was calculated by multiple t-tests in Prism6. For image analysis, significance value was calculated by Mann Whitney test in Prism6. P values under 0.05 were considered statistically significant and the following denotations were used: ***, P<0.001; **, P<0.01; * P<0.05.

## Acknowledgments

We thank F. Chisari for the gift of Huh7.5.1 cells; J. Zhong for mouse anti-NS5A antibody; S. Lemon, Z. Feng for providing HAV infectious clone and HM175/p16 virus; and T. Wakita, C. M. Rice, J. Bukh, and R. Bartenschlager for providing HCV strains; B. Qian for supporting software of imaging analysis. We apologize to many respected colleagues whom we could not cite because of space limitations.

Conceptualization: G.L., W.J.

Data Curation and Analysis: W.J.

Provision of materials: J.G.

Performing the experiment: W.J., P.M., F.M.

Project Administration and Supervision: G.L.

Original Draft Preparation: W.J.

Review & Editing: G.L..

All authors disclose no conflict of interest.

---

S1 Fig. An IPS-1-based reporter system for dual detection of HAV and HCV support robust HAV/HCV replication. (A) (B) Huh-7.5.1-GA/RC cell supports HAV and HCV infection as robustly as its progenitor Huh-7.5.1 cell, reflected by Western blot analysis of viral protein expression. (C) Dual fluorescence reporter system could detect HCV and HAV infection simultaneously and distinctly. The cell line Huh-7.5.1-GA/RC could distinguish two viral infections by distinct nuclear fluorescent color, with HAV green and HCV red. (D) HCV persistent infection did not affect TIM-1 expression.

S2 Fig. Extracellular viral structural protein level in HAV superinfection. (A) Western blot analysis of HAV VP2. (B) Elisa test of HCV core. Data represent the mean of three independent assays.

S3 Fig. HAV-induced cell death can be alleviated by intracellular pre-existing HCV. (A) Apoptosis/necrosis was detected by double dying Huh-7.5.1 of AnnexinV-EGFP and PI(propidium iodide) 72 hps(left). Percentage of cells stained with AnnexinV-EGFP or PI (right). In one randomly selected image, (±)AnnexinV-EGFP and (±)PI cells were counted and the percentage was calculated by the division of total cell numbers (B) Relative cell proliferation in the condition of indicated virus infection. MTT assay was applied to determine cell quantities by OD570 absorbance. Results showed the ratio of indicated value to the HCV persistent infection at 4hps. Data represent the mean of three independent assays.

S4 Fig. (A) HAV and (B) HCV replication under low MOI(≈0.01) of HM175/18f superinfection. (C) (A) HAV and (B) HCV replication in a long-time HM175/p16 superinfection. Intracellular viral RNA was quantified by RT-qPCR. Data represent the mean of three independent assays.

S5 Fig. Ectopic over-expression of HCV structural proteins did not disturb HAV life cycle. (A) Cell over-expressing HCV structural proteins (including E1, E2, core) was confirmed by western blot and designated as Huh-7.5.1-CNS2. (B) Comparison of HAV viral protein expression among Huh-7.5.1, Huh-7.5.1-GFP, and Huh-7.5.1-CNS2 cells 48 hours post HAV infection. (C) Comparison of HAV extracellular infectivity among Huh-7.5.1-GFP and Huh-7.5.1-CNS2 cells 48 hours post HAV infection.

S6 Fig. Strand-specific RT-qPCR quantification of HAV and HCV (+) (-) RNA copies.

S1 Tab. Primers for plasmid cloning of fluorescent reporter system.

S2 Tab. Primers for qPCR, strand-specific RT-qPCR and cloning cDNA region of negative HAV/HCV 3’UTR.

