## supplementary files for "A visualizable hepatitis A virus and hepatitis C virus coinfection model *in vitro*: coexistence of two hepatic viruses under limited competition in viral RNA synthesis"

### striking image

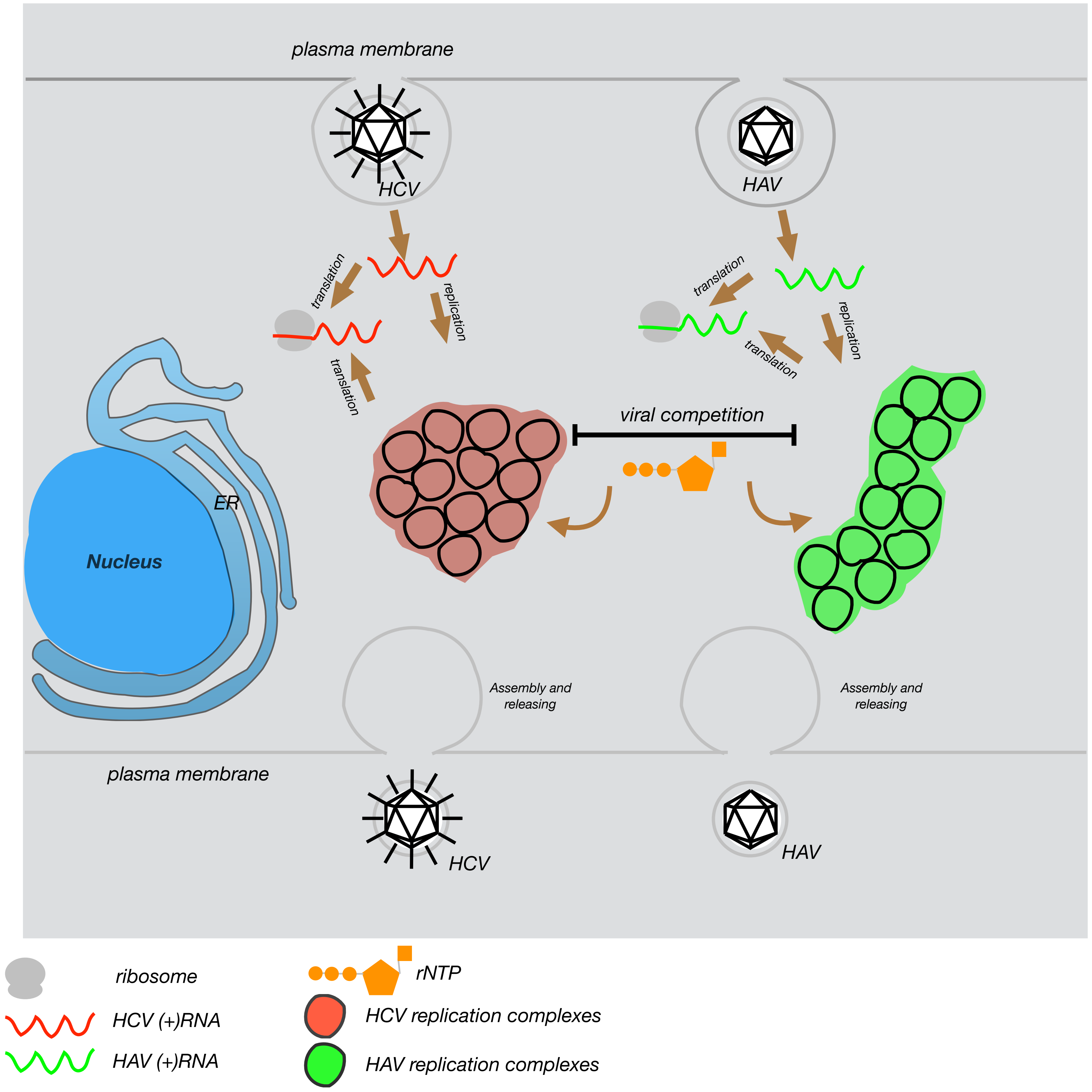

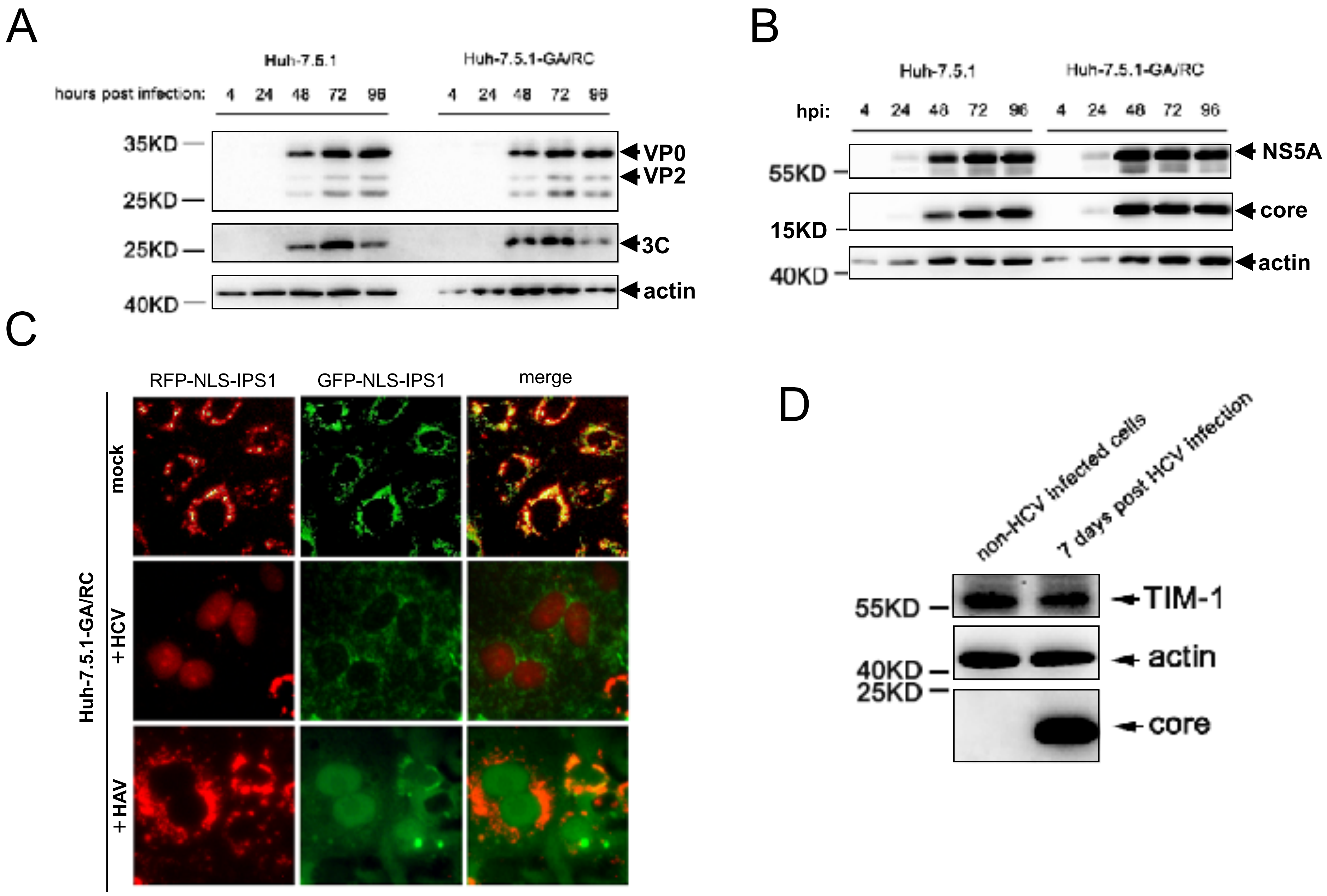
A
Huh-7.5.1
Huh-7.5.1-GA/RC
hours post infection: 4 24 48 72 96 4 24 48 72 96
35KD
25KD
25KD
40KD
VP0
VP2
3C
actin
B
Huh-7.5.1
Huh-7.5.1-GA/RC
hpi: 4 24 48 72 96 4 24 48 72 96
55KD
15KD
40KD
NS5A
core
actin
C
RFP-NLS-IPS1
GFP-NLS-IPS1
merge
Huh-7.5.1-GA/RC
mock
+HCV
+HAV
D
non-HCV infected cells
7 days post HCV infection
55KD
40KD
25KD
TIM-1
actin
core

sf1

A B

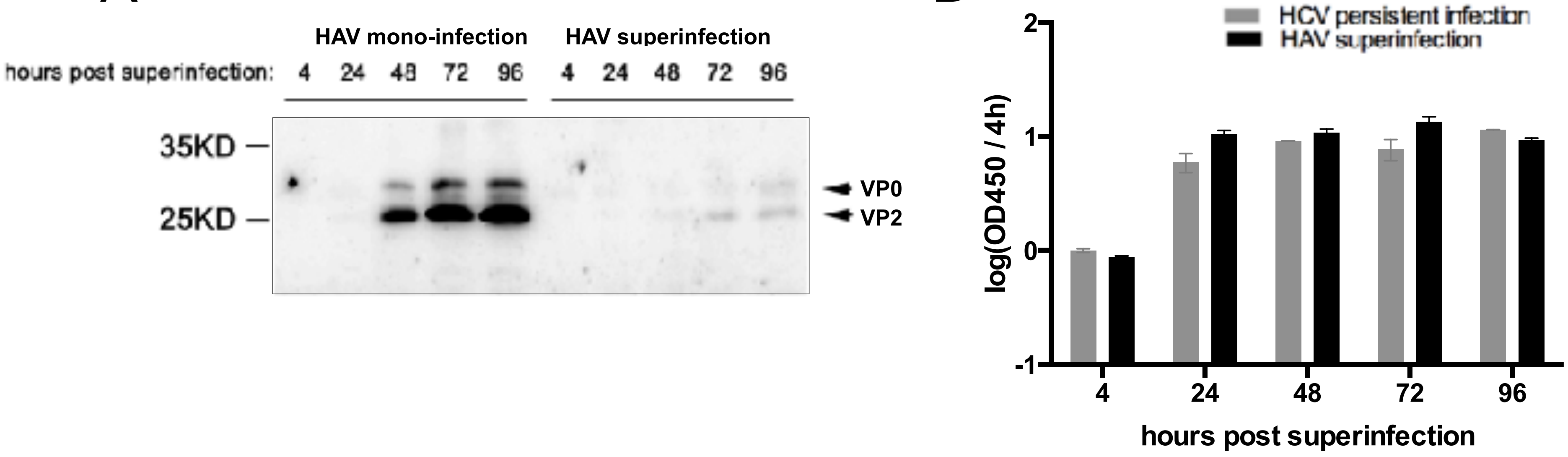
HAV mono-infection
HAV superinfection
hours post superinfection: 4 24 48 72 96 4 24 48 72 96
35KD
25KD
VP0
VP2
extracellular core Elisa
HCV persistent infection
HAV superinfection
log(OD450 / 4h)
2
1
0
-1
4
24
48
72
96
hours post superinfection

sf2

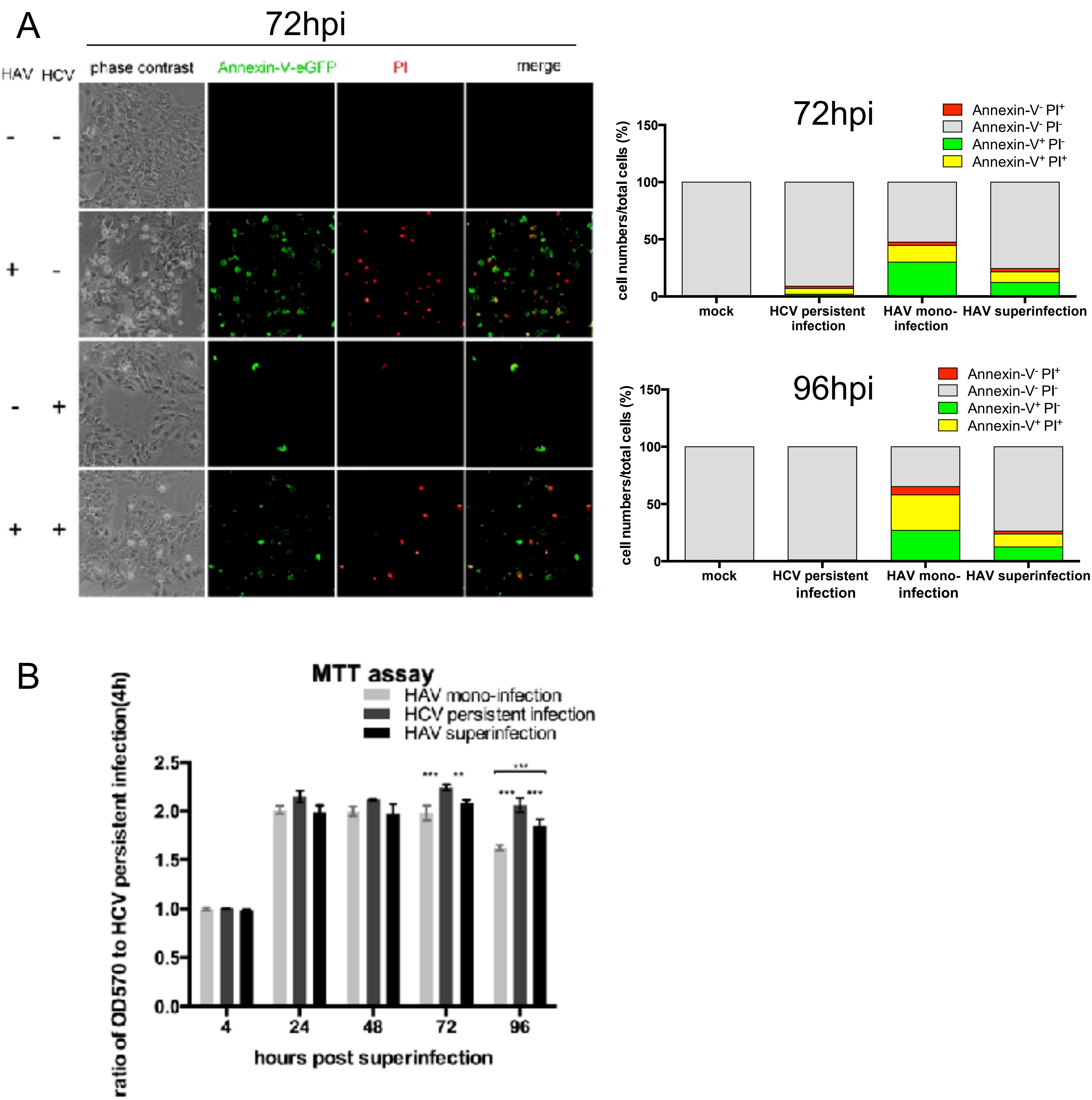
A
72hpi
HAV HCV
phase contrast
Annexin-V-eGFP
PI
merge
72hpi
96hpi
Annexin-V- PI+
Annexin-V- PI-
Annexin-V+ PI-
Annexin-V+ PI+
cell numbers/total cells (%)
mock
HCV persistent infection
HAV mono-infection
HAV superinfection
B
MTT assay
HAV mono-infection
HCV persistent infection
HAV superinfection
ratio of OD570 to HCV persistent infection(4h)
hours post superinfection

sf3

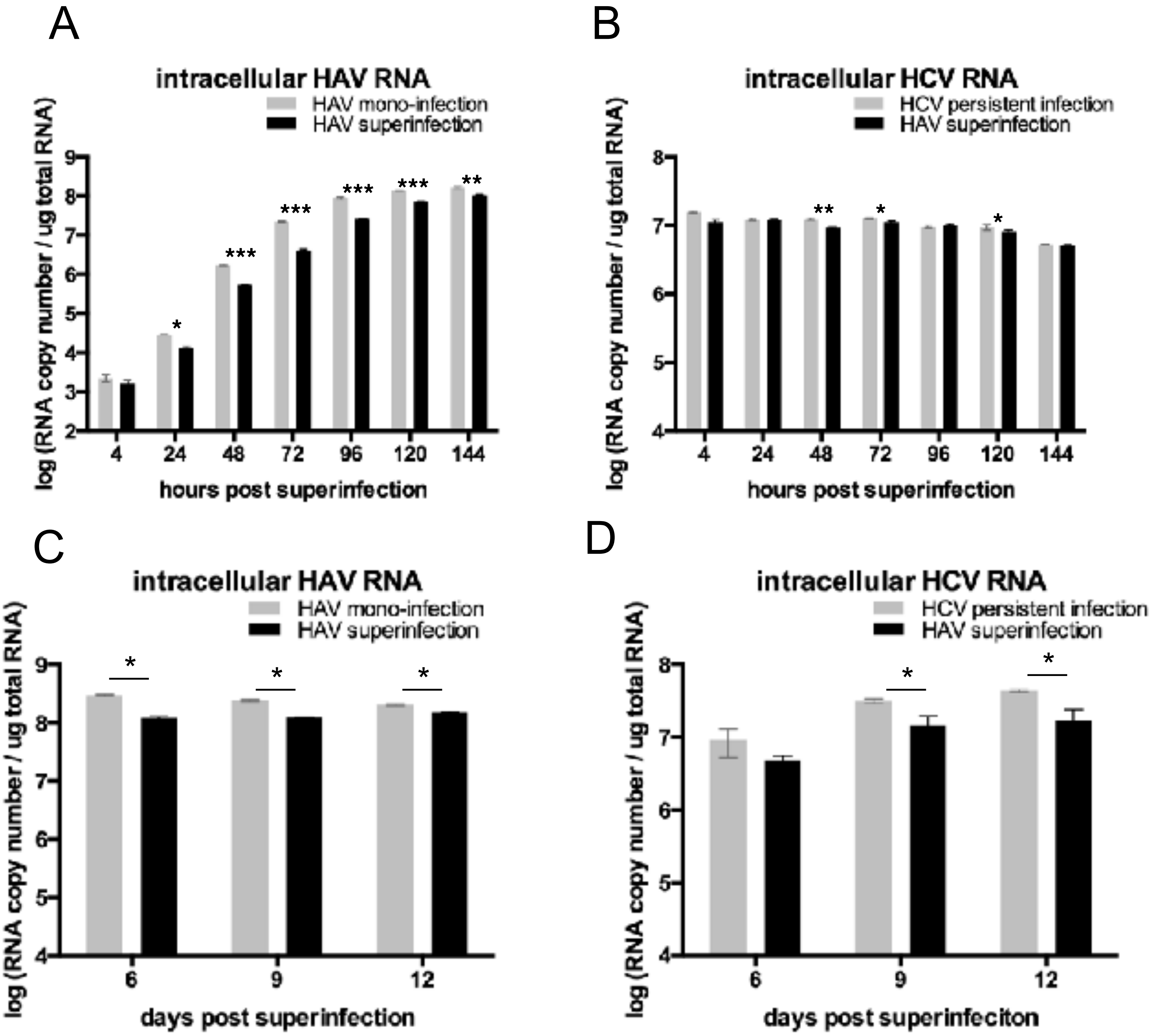
A
intracellular HAV RNA
HAV mono-infection
HAV superinfection
log (RNA copy number / ug total RNA)
hours post superinfection
B
intracellular HCV RNA
HCV persistent infection
HAV superinfection
log (RNA copy number / ug total RNA)
hours post superinfection
C
intracellular HAV RNA
HAV mono-infection
HAV superinfection
log (RNA copy number / ug total RNA)
days post superinfection
D
intracellular HCV RNA
HCV persistent infection
HAV superinfection
log (RNA copy number / ug total RNA)
days post superinfeciton

sf4

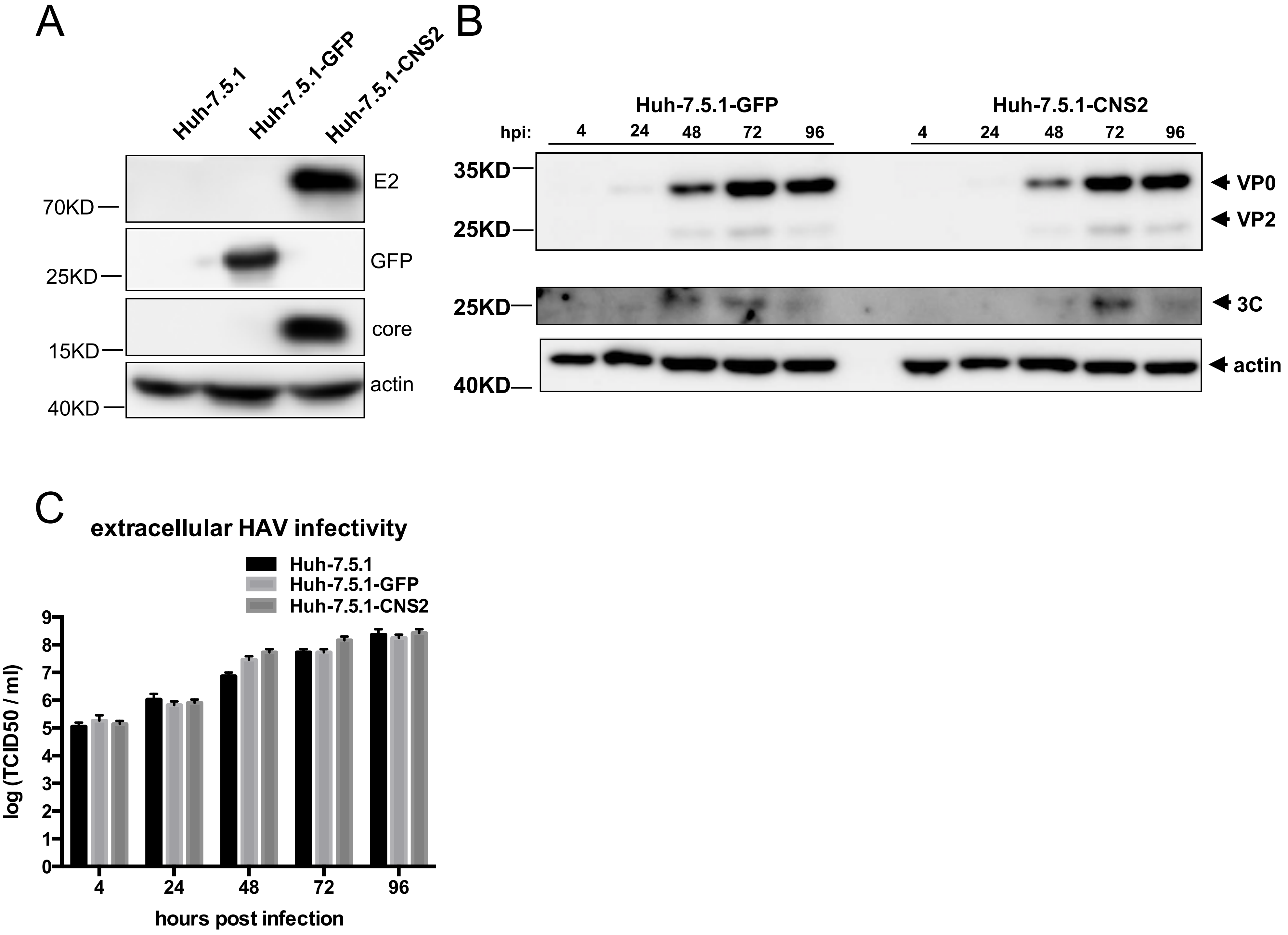

A
Huh-7.5.1
Huh-7.5.1-GFP
Huh-7.5.1-CNS2
E2
70KD
GFP
25KD
core
15KD
actin
40KD
B
Huh-7.5.1-GFP
Huh-7.5.1-CNS2
hpi: 4 24 48 72 96 4 24 48 72 96
35KD
25KD
VP0
VP2
25KD
3C
actin
40KD
C
extracellular HAV infectivity
Huh-7.5.1
Huh-7.5.1-GFP
Huh-7.5.1-CNS2
log (TCID50 / ml)
0 1 2 3 4 5 6 7 8 9
4 24 48 72 96
hours post infection


sf5

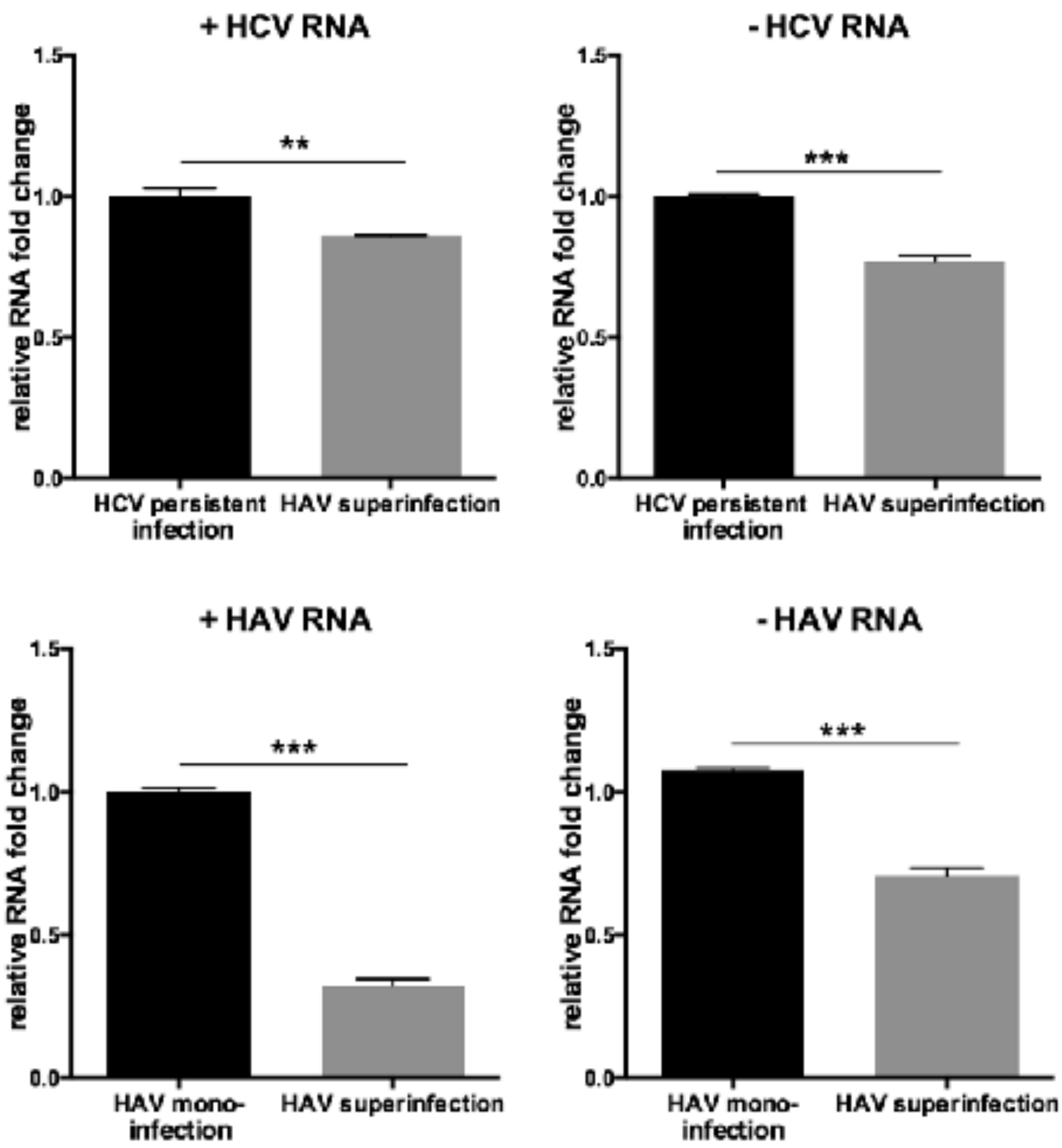
+ HCV RNA
- HCV RNA
+ HAV RNA
- HAV RNA
relative RNA fold change
1.5
1.0
0.5
0.0
**
***
HCV persistent infection
HAV superinfection
HAV mono-infection

sf6

### Table1

| **Primers for fluorescent reporter system** | |
|---|---|
| Primer name | 5'-3' sequence |
| eGFP-primer-F | ATACCTGCAGGATGGTGAGCAAGGGCGAGGAGC |
| NLS-IPS1(420)-R | **GATGCCAGCACGCCAGGCTT**AACTTTTCTCTTTTTCTTTGGATC |
| NLS-IPS1(420)-F | GATCCAAAGAAAAAGAGAAAAGTT**AAGCCTGGCGTGCTGGCATC** |
| eGFP-primer-R | AATACTAGTTTAGTGCAGACGCCGCCGGTACAGCACC |
| RFP-primer-F | TATCCTGCAGGATGAGCGAGCTGATC |
| NLS-IPS1(462)-R | AACTTTTCTCTTTTTCTTTGGATC |
| NLS-IPS1(462)-F1 | TCC***CGG***GGCACCTTTGGGATC |
| NLS-IPS1(462)-F2 | **GATCCAAAGAAAAAGAGAAAAGTT**TCC***CGG***GGCACCTTTGGGATC |
| RFP-primer-R | TATACTAGTTTAGTGCAGACGCCGC |
| Notes: The bold text represents NLS sequence for overlapping PCR; the underlined text represents the sequence of Sse8387I and SpeI for inserting PCR fragment into lentiviral plasmid pWPI-blr and pWPI-puro. | |

### Table2

| **qPCR primers** | **5'-3' sequence** |
|---|---|
| HAV-Forward | GGTAGGCTACGGGTGAAAC |
| HAV-Reverse | AACAACTCACCAATATCCGC |
| HCV-Forward | TCTGCGGAACCGGTGAGTA |
| HCV-reverse | TCAGGCAGTACCACAAGGC |
| **RT-primers** | **5'-3' sequence** |
| RT-HAV plus-RNA | GCTAGCTTCAGCTAGGCATCAACAACTCACCAATATCCGC |
| RT-HAV minus-RNA | GGCCGTCATGGTGGCGAATGGTAGGCTACGGGTGAAAC |
| RT-HCV plus-RNA | GGCCGTCATGGTGGCGAATTCTGCGGAACCGGTGAGTA |
| RT-HCV minus-RNA | GCTAGCTTCAGCTAGGCATCTCAGGCAGTACCACAAGGC |
| **tag-primers** | **5'-3' sequence** |
| tag-primer-F | GCTAGCTTCAGCTAGGCATC |
| tag-primer-R | GGCCGTCATGGTGGCGAAT |
| **pAC-minus(5'-3' sequence)** | |
| **gctagc**tcaggcagtaccacaaggcctttcgcaacccaacgctactcgggtagcagtcttgcgggggcacgcccaaatggccgggcatagagtgggtttatccaagaaaggacccagtcttcccggcaattccggtgtactcaccggttccgcagaccactatggctctcccgaacaactcaccaatatccgccgctgttaccctatccaaggcatctcttcatagaagtattagcctaagaggtttcacccgtagcctacc**aagctt** (268bp) | |
| Notes: Red: HCV qPCR region ; green:HAV qPCR region; underlined: primer targeted seuqences ; **bold:** restriction site of HheI or HindIII | |
